# Early-life stress alters chromatin modifications in VTA to prime stress sensitivity

**DOI:** 10.1101/2024.03.14.584631

**Authors:** Luke T. Geiger, Julie-Anne Balouek, Mason R. Barrett, Jeremy M. Thompson, Lisa Z. Fang, Lorna A. Farrelly, Andy S. Chen, Megan Tang, Shannon N. Bennett, Benjamin A. Garcia, Ian Maze, Meaghan C. Creed, Catherine Jensen Peña

## Abstract

Early-life stress increases sensitivity to subsequent stress, which has been observed at behavioral, neural activity, and gene expression levels. However, the molecular mechanisms underlying such long-lasting sensitivity are poorly understood. We tested the hypothesis that persistent changes in transcription and transcriptional potential were maintained at the level of the epigenome, through changes in chromatin. We used a combination of bottom-up mass spectrometry, viral-mediated epigenome-editing, RNA-sequencing, patch clamp electrophysiology of dopamine neurons, and behavioral quantification in a mouse model of early-life stress, focusing on the ventral tegmental area (VTA), a dopaminergic brain region critically implicated in motivation, reward learning, stress response, and mood and drug disorders. We found that early-life stress alters histone dynamics in VTA, including enrichment of histone-3 lysine-4 monomethylation — associated with open chromatin and primed or active enhancers — and the H3K4 monomethylase *Setd7*. Mimicking early-life stress through postnatal overexpression of *Setd7* and enrichment of H3K4me1 in VTA sensitizes transcriptional, physiological, and behavioral response to adult stress. These findings link early-life stress experience to long-term stress hypersensitivity within the brain’s dopaminergic circuitry, providing a mechanism by which early-life stress increases risk for mood and anxiety disorders later in life.

## INTRODUCTION

Early-life stress is a risk factor for mental health and substance use disorders, due to increasing sensitivity to subsequent stressors. (*1–4*). Mouse models of stress across the lifespan faithfully recapitulate these clinical findings, such that small behavioral differences after early-life stress are exaggerated in response to additional stress later in life (*5–7*). The ventral tegmental area (VTA) is a key dopaminergic brain region and has been extensively implicated in the pathophysiology of mood, anxiety, and substance-use disorders (*8, 9*). Stress-induced adaptations in gene expression and cellular activity in the VTA have been causally linked to changes in motivation, reward learning, and stress response (*10–14*). Indeed, ELS exerts prominent and long-lasting effects within the VTA and its projection targets in clinical and preclinical studies (*15–22*) including long-lasting alterations in transcription and epigenetic programming (*5, 6, 23–27*). We and others have observed both outright gene expression differences in VTA and other mesocorticolimbic brain regions following ELS, and latent gene expression changes that were revealed by a second hit of stress in adulthood (*5, 6, 28*). However, we lack a mechanistic understanding of how such latent gene expression responses are maintained, and how these molecular changes give rise to behavioral stress sensitivity. We hypothesize that these persistent changes in transcription and transcriptional potential in VTA are maintained at the level of the epigenome through changes in chromatin.

By dynamically responding to developmental and environmental cues, chromatin acts as a substrate of molecular memory in cells and facilitates adaptive gene expression responses to recurring stimuli, a phenomenon known as epigenetic priming (*29–33*). Post-translational histone modifications (PTHMs), including methylation, acetylation, OGlyc-NAc, serotonylation, and dopaminylation have been implicated in psychiatric disease and are altered in response to acute and chronic stress across the lifespan (*24, 27, 34–45*). However, the extent to which ELS alters the chromatin landscape of the VTA and whether such epigenetic changes prime response to adult stress as a mechanism of stress hypersensitivity are unknown. We used bottom-up mass spectrometry to characterize over 200 individual and combinatorial PTHMs in adult VTA following ELS. We then determined whether changes in PTHMs could be driven by changes in corresponding histone-modifying enzymes, focusing on the histone-3 lysine-4 monomethyltransferase *Setd7*. Finally, we found that overexpressing *Setd7* to drive elevated levels of H3K4me1 during development was sufficient to prime sensitivity to future stress at gene expression, physiological, and behavioral levels, effectively phenocopying ELS. Together, this establishes a chain of causality between ELS, H3K4me1-mediated chromatin remodeling, physiological adaptations in VTA, and increased susceptibility to stress in adulthood.

## RESULTS

### ELS alters histone methylation and acetylation dynamics in the VTA

We quantified proportions of PTHMs using unbiased, label-free mass spectrometry (LC-MS/MS) in VTA from adult male standard-reared and ELS mice (n= 3 replicates per group) (**Figure 1A**). Such analyses allow for simultaneous quantification of hundreds of histone PTHM states within the same sample, both in isolation and combinatorially with adjacent post-translational modifications; co-occurring combinations of modifications within a fragment are possible when that fragment includes multiple possible peptides that can be methylated or acetylated. We identified a unique pattern of single and combinatorial states of methylation and acetylation occurring on histone H3, histone H4 and H2A variant histone proteins following ELS relative to controls.

**Figure 1:**
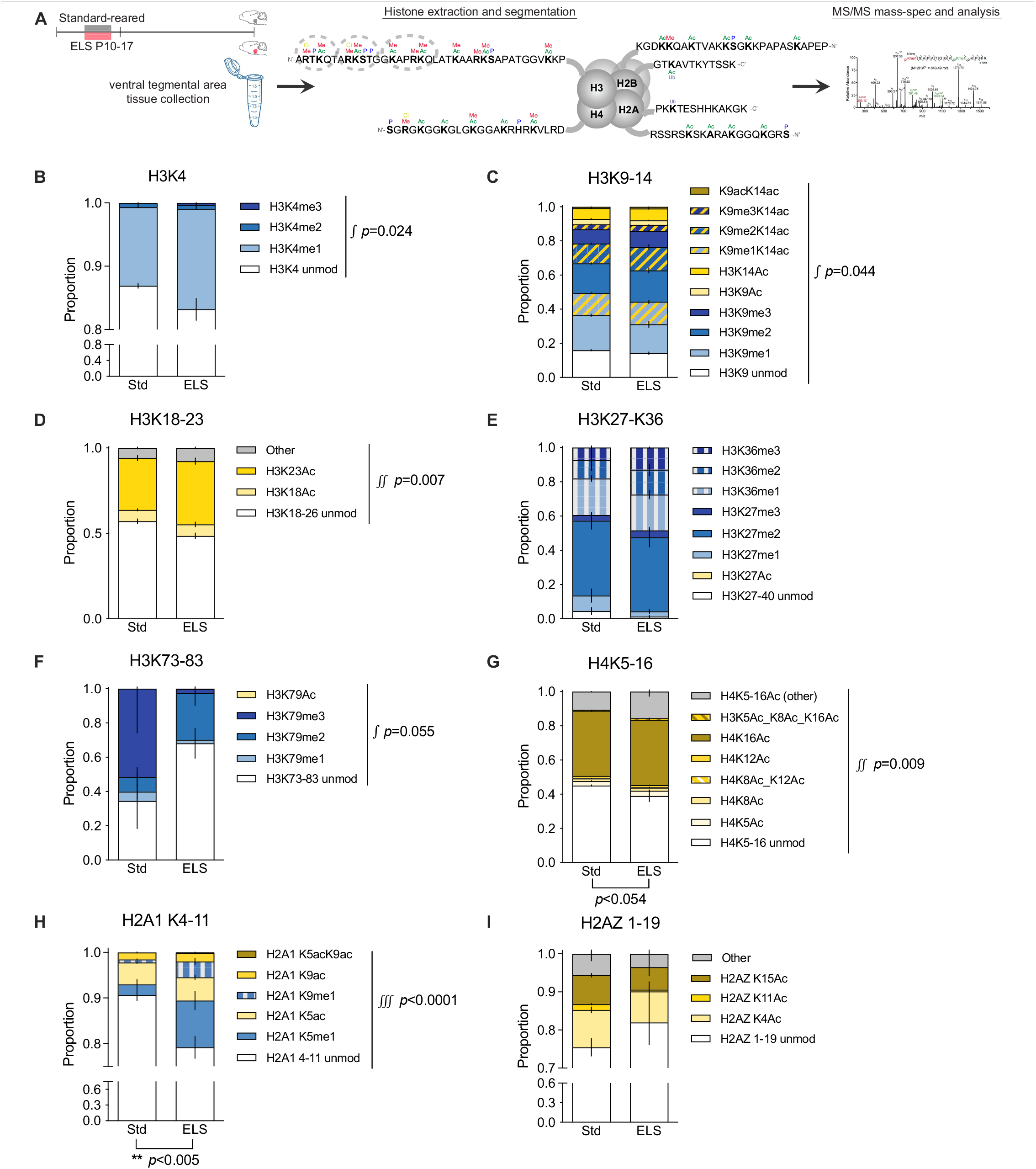
ELS alters proportions of histone modifications in VTA. **(A)** Study design overview: Male mice were either standard-reared (Std) or experienced ELS from P10-17, followed by standard conditions until adulthood. VTA was extracted, histones purified and histone tails fragmented, and subjected to LC-MS/MS. **(B-I)** Proportions of PTHM’s on each histone tail fragment for Std and ELS. Methylation is shown with blue shades, acetylation is shown with yellows, and dual modifications are shown with stripes. Vertical bars depict standard error from three biological replicates. There was an interaction between rearing condition and PTHM proportion for H3K3-8 [repeated measures ANOVA on arcsin-transformed ratios: F(3,12)=4.560, *p*=0.024]; H3K9-17 [F(9,36)=2.218, *p*=0.044]; H3K18-26 [F(3,12)=6.488, *p*=0.007]; H3K73-83 [F(4, 16)=2.912, *p*=0.055, trend-level]; H4K5-16 [F(7,28)=3.442, *p*=0.009]; and H2A1K4-11 [F(5,20)=15.53, *p*<0.0001]. There was a main effect of ELS within H2A1K4-11 [F(1,4)=31.02, *p*=0.005] and H4K5-16 [F(1,4)=7.30, *p*=0.054, trend level]. H2AZK1-9 also shown. Interactions between ELS and PTHM proportions are indicated by ∫ with the p-value shown; main effects of ELS are indicated below the bars.

Of the 27 peptide fragments detected, there was an interaction between rearing condition and PTHM proportion within six peptide fragments (**Figure 1B-I**): H3K3-8, H3K9-14, H3K18-26, H3K73-83 (moderate), H4K5-16, and H2A1K4-11. We also detected an effect of ELS within the H2A1K4-11 fragment (**Figure 1H**). There were no significant effects of ELS on methylation status of H3K27-K36, which has a prominent repressive role, and no H3K27Ac detected (**Figure 1E**). Within these fragments and H2AZ1-9 (**Figure 1I**), ELS increased proportions of fourteen PTHMs/combinations with large effect size (Cohen’s |*d*|>1.3) including: H3K4me1, H3K4me3, H3K9me3, H3K9me3-K14Ac, H3K9Ac-K14Ac, H3K23Ac, H3K36me2, H3K79me2, H4K8Ac, H4K8Ac-K12Ac, H4K5Ac-K8Ac-K16Ac, H2A.1K5me1, H2A.1K9me1, and H2A.1K5Ac-K9Ac (**Supplemental Figure 1**). ELS decreased proportions of unmodified H3K4, unmodified H3K18-26, unmodified H2A.1K4-11, and H2AZK11Ac (**Supplemental Figure 1**). A majority of these modifications are associated with open, active, primed or poised chromatin state, such that ELS would be predicted to increase transcriptional potential or facilitate gene expression responses. These results indicate that ELS alters histone dynamics in VTA that endure into adulthood.

### ELS alters the H3K4 monomethyltransferase Setd7

In order to determine whether changes in PTHMs could be driven by changes in corresponding histone-modifying enzymes, we examined published gene expression data from adult male VTA after ELS (**Figure 2A**; GEO accession GSE89692; (*5*)), and next validated expression levels of several enzymes in an independent cohort including both male and female mice exposed to ELS. From RNA-seq data, we examined normalized expression of four H3K4 methyltransferases (*Kmt2a, Kmt2b, Kmt2c*, and *Setd7*), two lysine demethylases (*Kdm1a* and *Kdm1b*), two H3K9 methyltransferases (*Setdb1* and *Ehmt2* also known as G9a), the H3K27 methyltransferase *Ezh2*, two histone acetyltransferases (*Crebbp* and *Ep300* also known as CBP/ p300), and five histone deacetylases (*Hdac2, Hdca3, Hdac5, Hdac11*, and *Sirt1*). The largest effect was for ELS to increase *Setd7* (**Figure 2A**).

**Figure 2:**
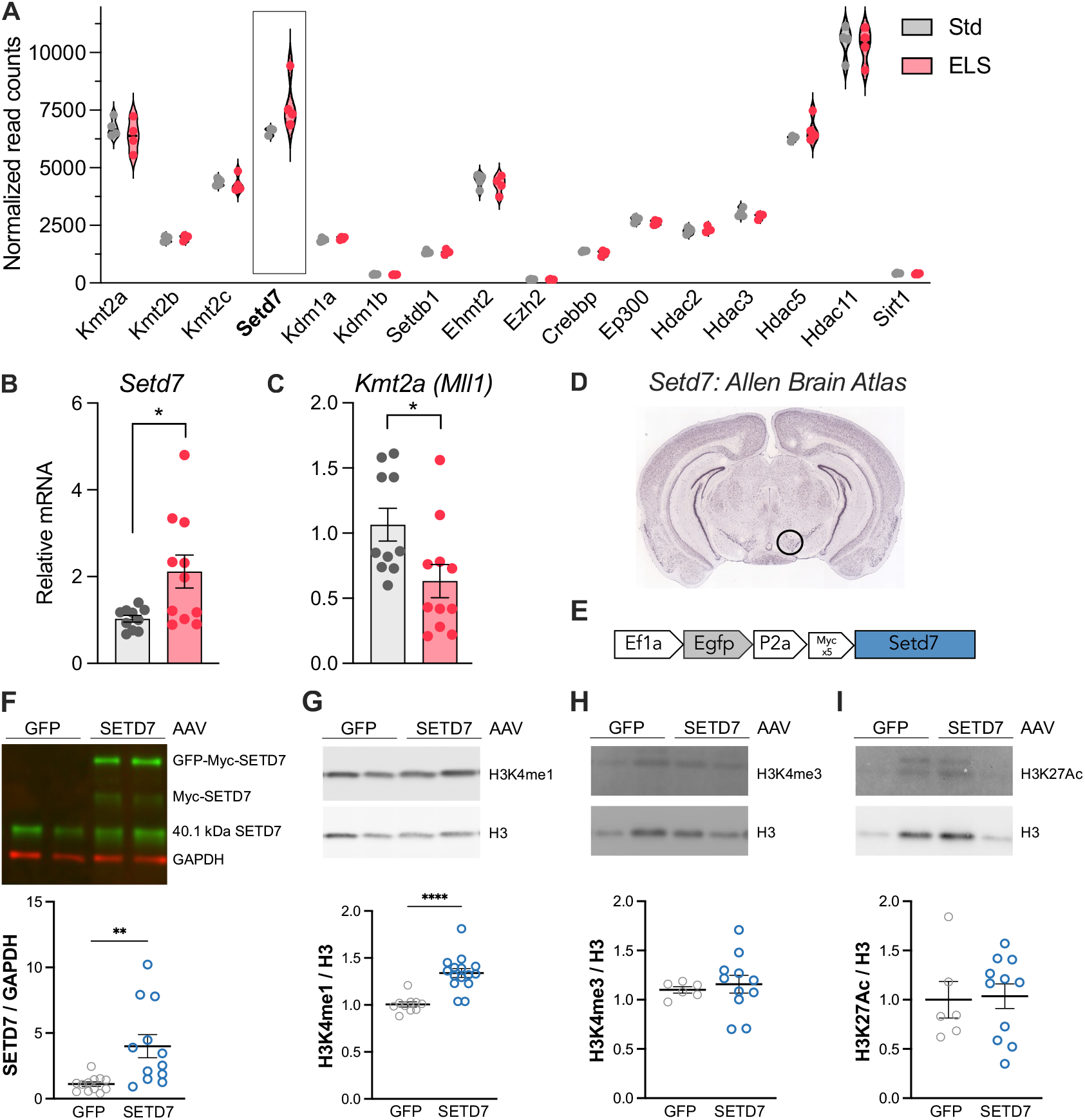
ELS increases the H3K4 monomethyltransferase *Setd7*. **(A)** Normalized expression (RPM) of histone modifying enzymes from published bulk RNA-sequencing data (GEO GSE89692) from the VTA of male mice that were standard- or ELS-reared. *Setd7* was increased by ELS with a Cohen’s d effect size *d*=1.078. **(B)** ELS increased *Setd7* mRNA in VTA in a qPCR replication cohort of male and female VTA [*t*(1,19)=2.656, *p*=0.0156]. **(C)** ELS decreased *Kmt2a* (MLL1) mRNA in a qPCR replication cohort of male and female VTA [*t*(1,19)=2.423, *p*=0.026]. **(D)** Allen Brain Atlas in situ hybridization shows that Setd7 is highly expressed in the dopaminergic VTA and substantia nigra at P56 (VTA is circled). **(E)** Viral vector design for overexpression of AAV8-Gfp-Myc-Setd7. **(F)** Bilateral AAV-Std7 overexpression in VTA increases total SETD7 protein 4-fold compared to AAV-Gfp injected control mice, including SETD7 protein at native 40.1 kDA size and SETD7 that remained fused to Myc and GFP tags [*t*(22)=3.211, *p*=0.004]. **(G)** *Setd7* overexpression increases H3K4me1 levels 34% relative to AAV-Gfp controls [*t*(22)=5.314, *p*<0.0001]. **(H)** H3K4me3 and **(I)** H3K27Ac levels remain unchanged. Example Western blots for protein or post-translational modifications are shown with either GAPDH or total histone H3 loading controls. Data are represented as mean ± SEM with \*\**p*<0.01 and \*\*\*\**p*<0.0001.

We validated that ELS increases *Setd7* expression in VTA within a qPCR replication cohort including both males and females (**Figure 2B**; n=10-11/group). In this cohort, ELS also suppressed the mixed-lineage leukemia (MLL)-type H3K4 methyltransferase *Kmt2a* (MLL1) (**Figure 2C**). In support of a role for *Setd7* in VTA, there is high *Setd7* expression within VTA and the nearby substantia nigra in the Allen Brain Atlas at P56 (**Figure 2D**; (*46*). Together, these findings support the view that the increase in H3K4me1 in VTA observed by mass spectrometry is attributable to increased expression of *Setd7*.

### Setd7 over-expression in mice specifically increases H3K4me1

SETD7 is a methyltransferase that specifically catalyzes the monomethylation of H3K4, without further catalyzing di- and tri-methylation at H3K4, or methylation at other histone lysine residues (*47, 48*). Given the interaction between ELS and H3K4 methylation state and increased expression of *Setd7* in VTA, we hypothesized that elevated H3Kme1 in VTA may facilitate the response to adult stress. To increase levels of H3K4me1 in VTA to test this hypothesis, we generated a *Setd7* overexpression vector (pAAV-EF1a-EGFP-P2A-Myc-Setd7-WPRE-hGHpA; **Figure 2E**) and corresponding e*Gfp* control vector without *Setd7* and packaged them into AAV8. We first validated that *Setd7* overexpression led to an increase in total SETD7 protein (**Figure 2F**) and H3K4me1 (**Figure 2G**) *in vivo*, in VTA. *Setd7* overexpression successfully enriched H3K4me1 levels by 34% relative to GFP control, normalized to total H3, consistent with its role as a monomethyltransferase (**Figure 2G**). However, there was no change in H3K4me3 or H3K27Ac (**Figure 2H-I**). These results indicate that *Setd7* overexpression specifically monomethylates H3K4 without recruiting further H3K4 methylation or other histone modifications associated with active gene expression.

### Juvenile Setd7 overexpression primes transcriptional response to adult stress

To test the hypothesis that SETD7-driven H3K4me1 enrichment in VTA enhances stress sensitivity, we overexpressed AAV-*Setd7* or AAV-*Gfp* control in developing VTA of male and female mice and evaluated gene expression, physiological, and behavioral responses to adult stress. For transcriptional (**Figure 3A**) and behavioral testing, injections were made at P14 during the ELS sensitive period (*5, 7*) and adult stress consisted of a “non-discriminatory” variant of adult social defeat stress that was previously found to be an effective stressor in both male and female mice simultaneously (*49*). However, aggressor mice in these experiments were observed to attack less overall with females present, and thus defeat was considered mild for these experiments compared to the standard ten days of male-only social defeat stress (e.g. (*5*). We also confirmed that our viral targeting of developing VTA was successful and relatively specific (**Figure 3B**). This was a particular challenge, given the young age of pups during surgery and deep-brain location of the VTA.

**Figure 3:**
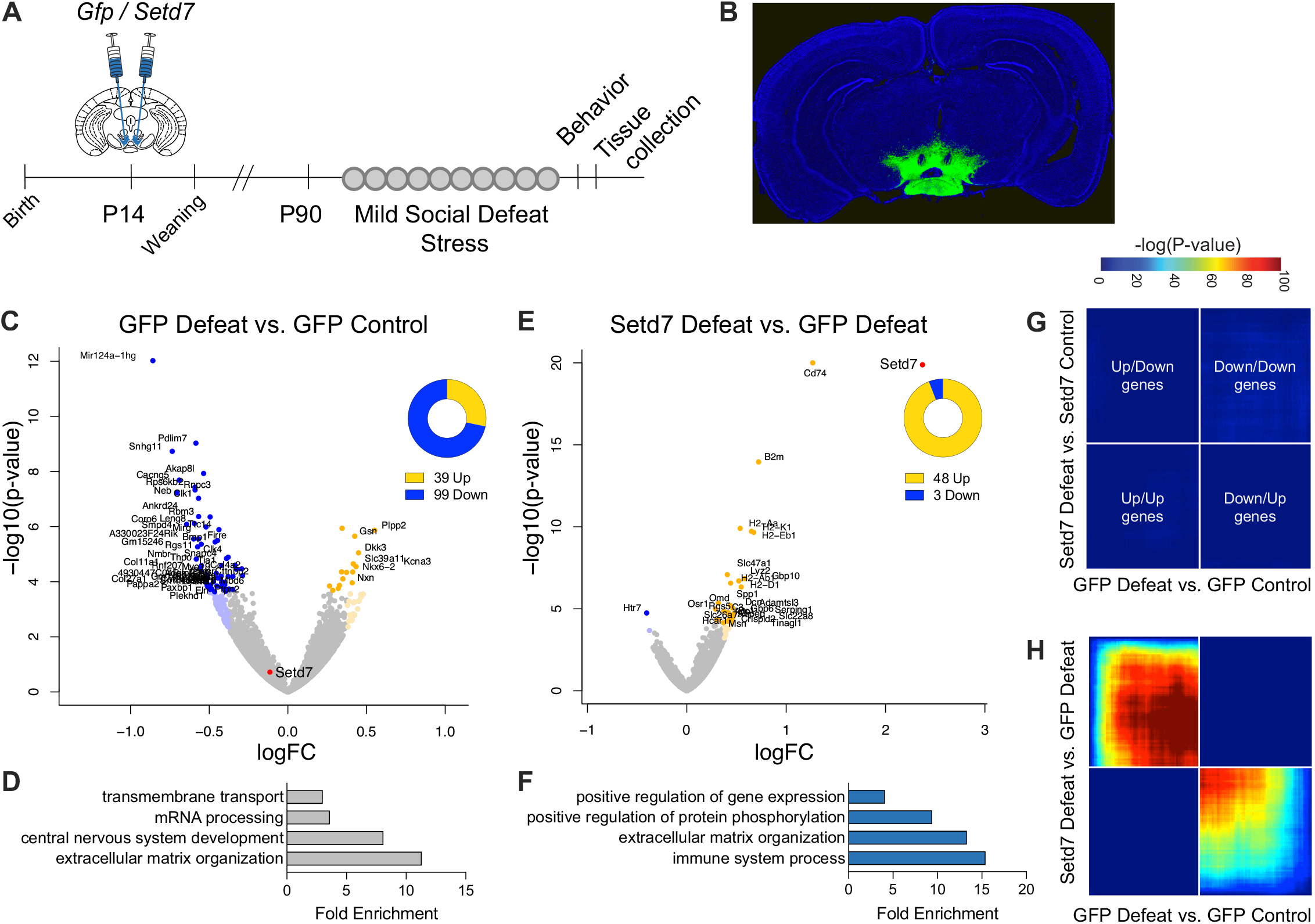
Juvenile *Setd7* overexpression augments transcriptional responses to adult stress. **(A)** Experimental design: AAV-Setd7 or AAV-Gfp was bilaterally injected into VTA of male and female mice at P14. VTA was collected two days after mild social defeat stress. **(B)** Example GFP confirmation of the spatial accuracy of juvenile AAV injections. **(C)** Volcano plot of differentially expressed genes (DEGs) in VTA among AAV-Gfp mice subjected to mild social defeat stress compared to control conditions, demonstrating 72% of DEGs (*p*.*adj*<0.1) are downregulated. Genes with *p*.*adj*<0.05 are shown in darker colors and additional genes with uncorrected *p*<0.01 and Log2FC<|0.375| are shown in lighter colors. **(D)** Gene ontology enrichment analysis of DEGs (*p*.*adj*<0.1) downregulated by GFP-Defeat vs GFP-Control. **(E)** Volcano plot of DEGs in VTA between stressed AAV-Setd7 mice compared to stressed AAV-Gfp mice, demonstrating that 94% of DEGs were upregulated. Setd7 is the most upregulated gene (Log2FC=2.37; *p*.*adj*=2.13×10-130, truncated 2.13×10-20 at for visualization). **(F)** Gene ontology enrichment analysis of DEGs upregulated by Setd7-Defeat vs Gfp-Defeat. Rank-rank hypergeometric overlap analysis comparing genome-wide similarities between **(G)** Setd7-Defeat vs Setd7-Control and Gfp-Defeat vs Gfp-Control comparisons, and **(H)** Setd7-Defeat vs Gfp-Defeat and Gfp-Defeat vs Gfp-Control comparisons. Genes are signed by Log2FC and ranked by p-value from most to least significantly regulated from the middle to outer corners; significance of overlap [−log10(p-value)] in a hypergeometric test is color-coded, with a fixed maximum of 100 across all comparisons.

Using bulk-tissue RNA-sequencing, we determined whether developmental overexpression of *Setd7* and accompanying H3K4me1 enrichment is associated with enhanced *transcriptional* reactivity to adult stress, as the primary consequence of epigenetic change is transcriptional. Because deposition of H3K4me1 by *Setd7* overexpression was assumed to be broad across the genome, we focused on pattern-level changes rather than individual genes. We first examined the impact of social defeat stress on gene expression within *Gfp* control mice and found that defeat stress predominantly decreased gene expression, with 99 genes down-regulated (72%) and 39 genes up-regulated (differential expression by DESeq2, *p*.*adj*<0.1) (**Figure 3C**). This pattern of largely suppressed gene expression is consistent with previous findings that approximately 70% of genes altered by adult stress were down-regulated in VTA (*6*). Genes down-regulated by defeat stress among *Gfp* control mice were enriched for functions related to extracellular matrix organization, central nervous system development, mRNA processing, and transmembrane transport (**Figure 3D**). We then examined the impact of stress on gene expression among mice epigenetically primed by *Setd7* vs *Gfp* and found that *Setd7* predominately increased gene expression responses to stress, with 48 genes up-regulated (94%) and only 3 down-regulated **(Figure 3E**). As expected, *Setd7* was the top most enriched gene (**Figure 3E**). Genes up-regulated by *Setd7* overexpression and defeat were enriched for functions related to positive regulation of gene expression, protein phosphorylation, extracellular matrix organization, and immune response (**Figure 3F**). We next asked whether the general decrease in gene expression following social defeat is reversed by *Setd7* overexpression. We found marked differences in the effect of social defeat stress within *Gfp* and *Setd7* groups across the genome using rank-rank hypergeometric overlap analysis (**Figure 3G**). However, the effect of stress within *Gfp* mice was largely reversed by *Setd7*, with genes normally down-regulated by stress predominately upregulated by *Setd7* (**Figure 3H**). These findings indicate that epigenetic priming within VTA by overexpression of *Setd7* increases positive transcriptional reactivity to adult stress.

### Juvenile Setd7 overexpression enhances dopamine neuron excitability after adult stress

Stress is known to alter excitability of VTA dopamine neurons, and these adaptations are causally related to behavioral susceptibility to stressors (*12, 13*). To test whether developmental overexpression of *Setd7* and H3K4me1 enrichment altered function of VTA dopamine neurons, we injected AAV-*Setd7* or AAV-*Gfp* in juvenile VTA (P21-24). Mice were aged into adulthood, before exposure to a mild unpredictable stress paradigm before performing patch clamp recordings of VTA dopamine neurons (**Figure 4A-B**). *Setd7* overexpression had no effect on excitability of dopamine neurons in control animals: there was no difference in the input-output curve (**Figure 4C**), action potential threshold (**Figure 4E**) or membrane input resistance (**Figure 4F**) between *Setd7* overexpression and *Gfp* controls. However, dopamine neuron excitability following adult stress was increased by *Setd7*, with greater action potential firing to increasing current injection as measured by increased area-under-curve (**Figure 4D**). Moreover, *Setd7* overexpression was associated with increased hyperpolarization-activated (*I*_h_) current, as measured by voltage sag in response to current injection (**Figure 4G**). This increase in depolarizing *I*_h_ current promotes activity of dopamine neurons by bringing the cell closer to action potential threshold during membrane hyperpolarizations, thereby facilitating neuronal firing (*50*). Enhanced *I*_h_ current following social defeat stress in adulthood has been associated with increased behavioral susceptibility (*13*). Together, these electrophysiology results establish that overexpression of *Setd7* alone does not alter the basal electrophysiological characteristics of dopamine neurons, but promotes cellular plasticity of dopamine neurons in response to stressful stimuli.

**Figure 4.**
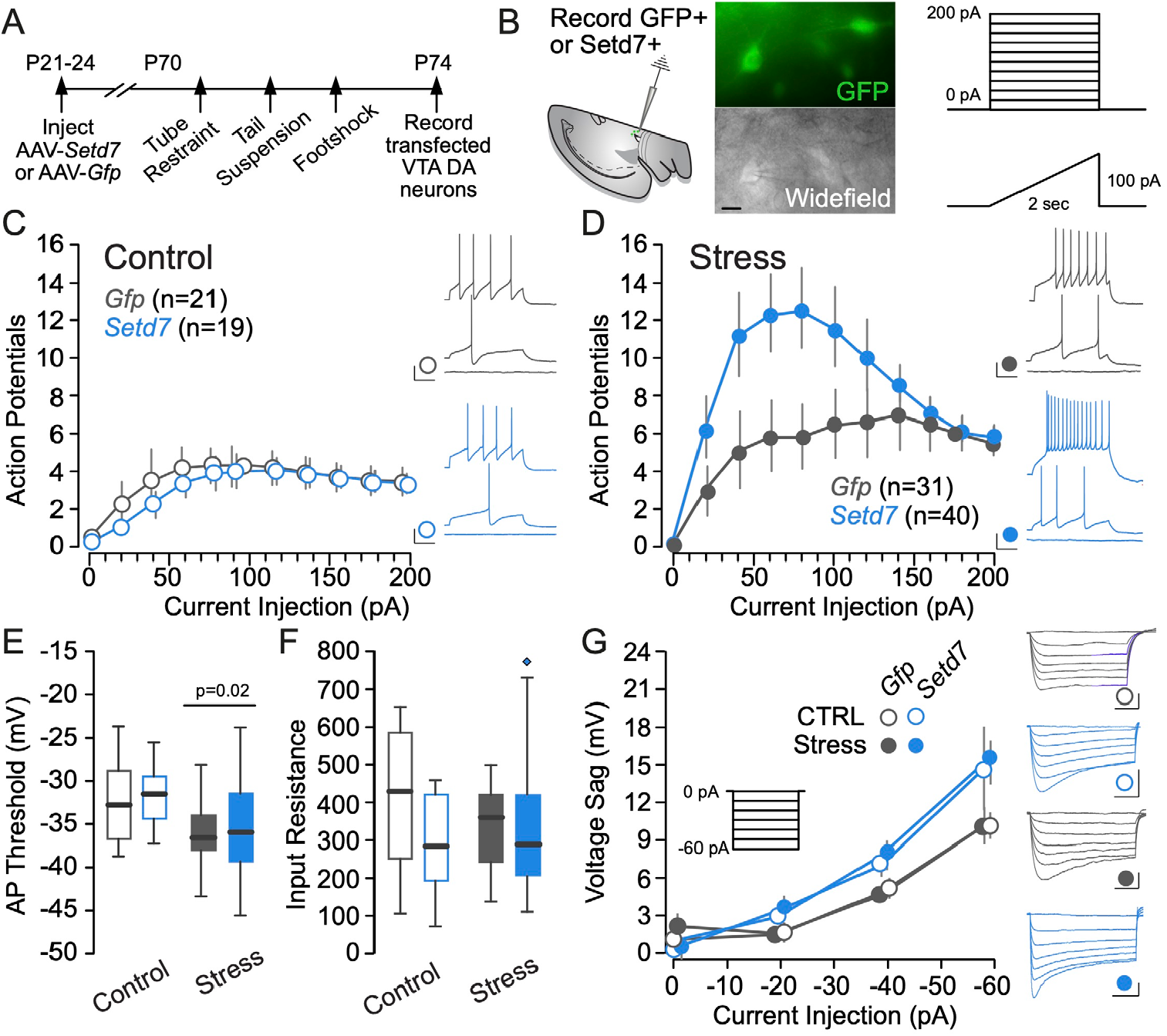
Juvenile *Setd7* overexpression enhances dopamine neuron excitability after adult stress. **(A)** Timeline of experiments: mice were transfected with AAV-*Setd7* or AAV-*Gfp* virus. After 4 weeks of expression, “Stress” mice underwent three days of distinct inescapable stressors, and patch clamp recordings were performed the subsequent day. **(B)** Schematic of recording and example image of viral transfection (right). Scale bar = 50 µm. **(C-D)** Input-output curve of action potentials generated in response to successive current injections in Control (C) and Stressed (D) mice. There was a significant effect of stress (F=6.70, *p*=0.011) and of *Setd7* (F=4.10, *p*=0.046) on excitability as reflected by the area under the input-output curve. **(E)** Stress had a significant effect of reducing AP threshold (CTRL-*Gfp*: 32.12 ± 1.22, CTRL-*Setd7*: 31.38 ± 0.84, Stress-*Gfp*: 35.93 ± 1.00, Stress-*Setd7*: 34.49 ± 1.12, F_Stress_=10.01, *p*=0.002). **(F)** There was no effect of stress or virus on input resistance (CTRL-*Gfp*: 370.51 ± 44.62, CTRL-*Setd7*: 349.01 ± 71.79, Stress-*Gfp*: 338.32 ± 25.25, Stress-*Setd7*: 305.27 ± 23.87). **(G)** *Setd7* expression significant increased peak *I*_h_ current, relative to *Gfp* (Normalized Peak Current *Setd7*: 17.23 ± 0.08 mV, *Gfp*: 11.3 ± 0.058 mV, *p*=0.023).

### Setd7 overexpression in VTA enhances behavioral sensitivity to adult stress

Finally, to determine whether developmental enrichment of H3K4me1 increases behavioral sensitivity to stress, mimicking ELS, we overexpressed *Setd7* or *Gfp* in male and female mice and used a within-subject testing design to assess for social and exploratory avoidance behaviors before (“pre”) and after (“post”) mild social defeat stress (**Figure 5A**). There was a main effect of mild social stress to decrease social interaction behavior, indicative of stress susceptibility in mice (*51*), and an interaction between *Setd7* and stress such that stress reduced social interaction behavior only among *Setd7* mice (**Figure 5B**). There was no effect of sex. Among *Setd7* but not *Gfp* mice, stress increased the proportion considered susceptible and decreased the proportion considered resilient (**Figure 5C**). Similarly, within an open field test, mild social defeat stress decreased center time among *Setd7* mice but not *Gfp* mice (**Figure 5D**). Together, these results indicate that developmental *Setd7* overexpression increases sensitivity to mild adult social defeat stress among both male and female mice.

**Figure 5:**
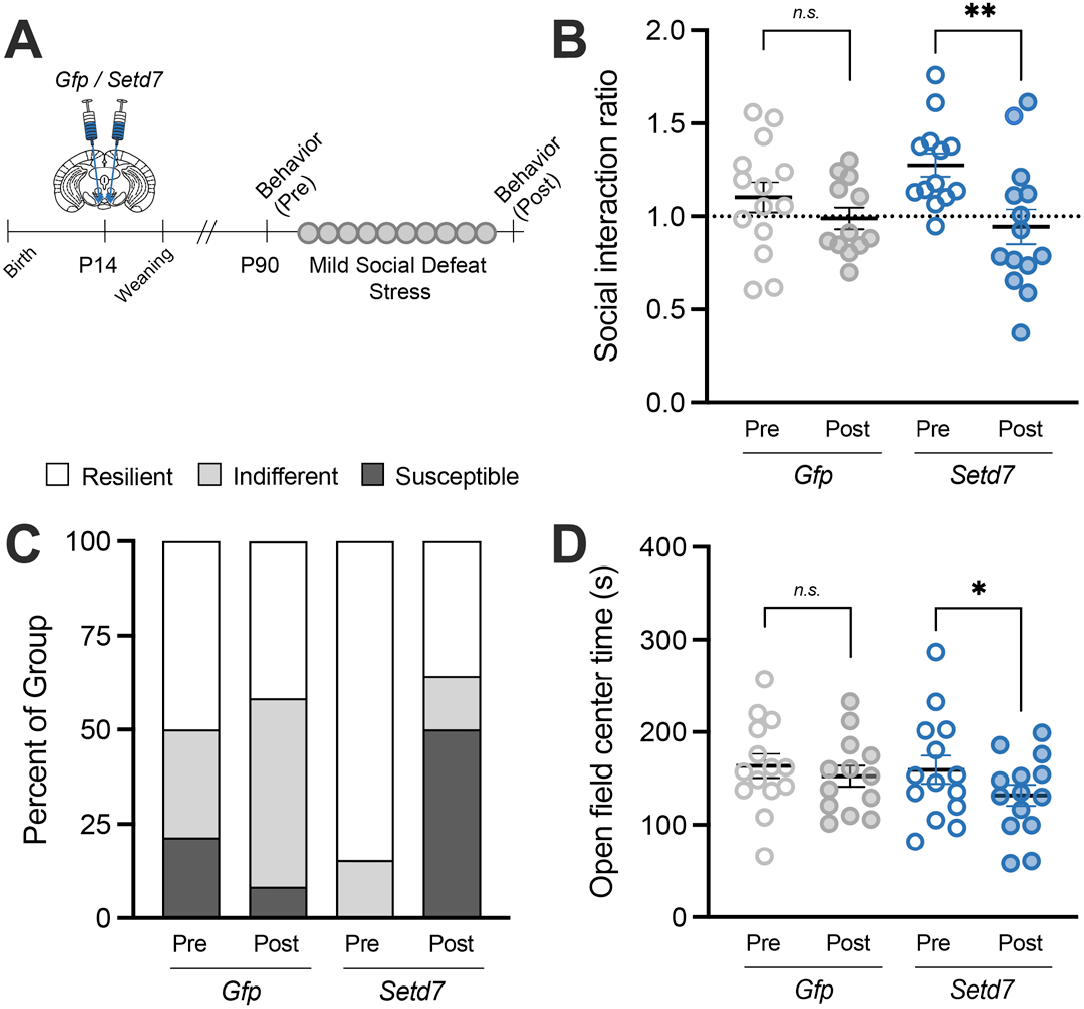
Juvenile Setd7 overexpression enhances behavioral sensitivity to adult stress. **(A)** Experimental design: AAV-*Setd7* or AAV-*Gfp* was bilaterally injected into VTA of male and female mice at P14. Behavior was quantified before (pre) and after (post) mild social defeat stress. **(B)** Social interaction ratio (time spent investigating a novel aggressive mouse vs empty enclosure) was reduced by mild social defeat stress among *Setd7*-overexpressing mice [main effect of social defeat: *F*(1,49)=8.240, *p*=0.0060; interaction between *Setd7* and defeat: *F*(1,49)=2.001, *p*=0.163; post-hoc test with Sidak multiple comparisons correction was significant for *Setd7* (*p*=0.0071) but not *Gfp* (*p*=0.5284)]. **(C)** 50% of *Setd7*-overexpressing mice were categorized as susceptible to mild social defeat stress, compared to 8.3% of *Gfp* mice (two-tailed two-proportion z-test: Z=2.96, *p*=0.003). Resilience to stress was also reduced by *Setd7* overexpression (Z=2.58, *p* =0.010). **(D)** Time spent in the center of an open field was reduced by mild social defeat stress [main effect of stress: two-way ANOVA with paired effects: *F*(1,25)=4.616, *p*=0.0416] among AAV-Setd7 (*p*=0.043) but not AAV-Gfp mice (*p*=0.395).

## DISCUSSION

ELS increases sensitivity to subsequent stress, which has been observed in both clinical (*2, 52, 53*) and preclinical studies (*5, 7, 54, 55*) at behavioral, cellular activity (*56*), and gene expression (*6, 28*) levels. We tested the hypothesis that persistent changes in transcription and transcriptional potential in VTA were maintained at the level of the epigenome through changes in chromatin. Here, using a combination of bottom-up mass spectrometry, viral-mediated epigenome-editing, RNA-sequencing, electrophysiology, and behavioral quantification, we show that ELS alters histone dynamics in VTA and that mimicking one such prominently altered histone modification augments transcriptional response to stress, enhances dopamine neuron excitability, and ultimately sensitizes behavioral response to stress.

ELS triggered enduring changes among a variety of histone methylation and acetylation modifications in VTA, across histones H3, H2A, and H4 (**Figure 1; Supplemental Figure 1**). Interestingly, 75% of histone modification changes were associated with a permissive (open, active, primed, or poised) state. This broad finding suggests that ELS may facilitate transcriptional response to future stress via a more permissive, or “primed,” chromatin state, although a direct quantification of chromatin accessibility in VTA following ELS is needed. These findings are consistent with broad and dynamic histone modifications reported following ELS in NAc, a direct target of VTA dopamine neurons, including an increase in H3K4me1 in adolescent males and females and adult males (*27*). Across histone H3 modifications often associated with gene repression, we found that ELS increases both H3K9me3 and H3K36me2. Recently, the co-occurrence of H3K9 and H3K36 methylation has been hypothesized to ‘bookmark’ enhancers for future transcriptional activity during embryonic stem cell differentiation (*57*), although it is not yet known whether the dual-occurrence of these marks functions similarly in the postnatal brain.

Our mass spectrometry data from ELS-treated mice is consistent with prior investigations of individual PTHMs. For example, H3K9Ac in VTA was reduced by a single 24-hour maternal separation on postnatal day P9 (*40*) and by ELS from P10-17 here. Interestingly, previous work found that reduced VTA H3K9Ac was associated with impaired GABAergic synaptic plasticity and increased long-term depression of GABAergic neurons onto dopamine neurons, resulting in metaplasticity and increased dopamine neuron excitability and dopamine release in the ventral striatum (*23, 26, 40*). Together, these findings indicate consistent and functional relevance of ELS-induced chromatin modifications in VTA, linking molecular changes with neuronal activity and behavior, as well as the need for cell-type specificity in future studies.

Despite changes among a variety of histone modifications in VTA, we found few enduring changes in the enzymes that write and erase these modifications. In fact, *Setd7* was the only significantly up-regulated histone-modifying enzyme (**Figure 2A-C**). SETD7 (SET7/9, KMT7) is a histone lysine methyltransferase that deposits a single methyl group at H3K4, often at gene promoters (*29, 47, 48, 58, 59*). Its role as a specific monomethyltransferase is unique relative to other common H3K4 methyltransferases (i.e., MLL1, MLL2, MLL3, MLL4, SET1A, and SET1B, encoded by *Kmt2a-f*, respectively) that are capable of serially adding di- and tri-methylation to H3K4, and in some cases have higher substrate specificity for methylated H3K4 (*60, 61*). The ability of *Setd7* overexpression to increase H3K4me1 but not H3K4me3 in our experiments **(Figure 2G-I)** corroborates its role as a selective monomethyltransferase. H3K4me1 and H3K4me3 have distinct enrichment across the genome, with H4K3me1 marking both active promoters and enhancers and poised/primed regulatory regions, while H3K4me3 is highly enriched at active promoter regions (*29*). These distinct distribution patterns require independent regulation themselves and serve different functions, suggesting specificity in the impact of ELS on genomic regulation.

Confirming the role of H3K4me1 in “priming” transcriptional responses, juvenile overexpression of *Setd7* in VTA increased transcriptional, physiological, and behavioral responses to adult stress (**Figures 3-5**). These findings demonstrate that *Setd7* overexpression in VTA is sufficient to recapitulate ELS-induced enhanced stress sensitivity. The increase in upregulated genes after adult stress with *Setd7* overexpression was notable in contrast to largely decreased VTA transcription in *Gfp* control mice. This decrease in transcription with social defeat is consistent with earlier work which also reported largely decreased transcription after adult stress in VTA (*6*). Among the most highly regulated genes are solute carrier protein family genes (**Supplemental Data**), which are critical for regulating excitability and function of dopaminergic neurons by maintaining chloride gradients and allowing for neurotransmitter packaging and release. However, we are cautious to put too much emphasis on individual genes altered by *Setd7* overexpression here given that this AAV overexpression vector does not allow us to control where H3K4me1 is deposited across the genome. It is likely to be deposited in genomic regions that are already open and accessible to the enzyme. This begs the need for future research with more refined tools such as CRISPR/dCas9-based targeted epigenetic modifications for improved specificity (*62–64*). Nevertheless, together, these findings are consistent with the interpretation that SETD7-driven H3K4me1 enrichment in VTA potentiates future transcriptional and neurophysiological responses to stress. This constitutes a potential mechanism by which ELS drives persistent increased sensitivity to stress experienced in adulthood.

*Setd7 is* enriched in the VTA and substantia nigra relative to surrounding regions (**Figure 2D**), suggesting a potential role in dopaminergic modulation, which was corroborated by our electrophysiology results. While *Setd7* overexpression alone had no significant effect on VTA dopamine neuron excitability, functional changes following mild stress were exaggerated in neurons overexpressing *Setd7* (**Figure 4D**). Stress has been shown to enhance firing activity of VTA dopamine neurons (*65*), which is causally related to behavioral adaptations to stressful stimuli (*13*). Consistent with this prior work, we observed increased firing to current injection after stress in both our *Gfp* control and *Setd7* groups. However, the excitability of VTA dopamine neurons was significantly greater in neurons overexpressing *Setd7*, suggesting the enhanced genomic reactivity mediated by *Setd7* was sufficient to increase functional sensitivity of VTA dopamine neurons to stressful stimuli.

While the current study demonstrates sufficiency of juvenile *Setd7* and H3K4me1 enrichment to prime stress sensitivity, we do not yet know whether these effects are specific to development, nor whether removing H3K4me1 or preventing its accumulation would render mice more resistant to stress. To this point, LSD1 (*Kdm1a*), a H3K4 demethylase, has been previously implicated in adult stress response. Adult stress (either social defeat or chronic variable stress) reduces LSD1 protein in hippocampus and *Kdm1a* mRNA in the medial prefrontal cortex (*66, 67*), potentially implicating regulation of H3K4 methylation across the brain. Reduction of *Kdm1a* reduced immediate early gene (*Egr1, Fos*) levels via reduced H3K4 methylation at their promoters, alleviated hypersensitivity to pain, and reduced stress-associated behaviors in mice (*66, 67*). Together with our current work, this reveals a critical role for regulation of H3K4 methylation in stress response.

A new question inspired by our findings is whether H3K4me1 plays a role in increased sensitivity to *non-stressful* stimuli, such as stimuli with positive valence. If chromatin priming of the VTA with *Setd7* and H3K4me1 drives sensitivity to aversive experiences in adulthood, it is possible that the VTA could equally become hyper-responsive to non-stressful environmental conditions, such as environmental enrichment. Activity of discrete populations of VTA dopamine neurons encodes stimuli of opposing valence, including both unexpected aversive and unexpected rewarding experiences (*68, 69*). The potential for epigenetic priming to drive sensitivity to positive, enriching experiences would open new possible avenues for intervention for individuals who have experienced childhood adversity.

In conclusion, enduring changes in histone modifications suggest that ELS facilitates response to future stimuli by promoting a more permissive chromatin state in the VTA. Mimicking ELS by increasing the monomethyltransferase *Setd7* and H3K4me1 promotes transcriptional, functional, and behavioral hypersensitivity to future stress. Together these findings establish epigenetic priming within the VTA as a biological mechanism through which ELS programs lifelong stress sensitivity.

## ACKNOWLEDGEMENTS

Thank you to Eric Nestler at Mount Sinai for support while generating tissue for mass-spec experiments and comments on an earlier version of the manuscript; to Natarajan Bhanu in the Garcia Lab for help processing mass-spec samples; to Esteban Engel and the Princeton Neuroscience Viral Core for assistance generating the AAV-*Setd7* overexpression vector; and to Wei Wang and the Genomics Core Facility in the Lewis-Sigler Institute for Integrative Genomics at Princeton where RNA-seq was performed.

## AUTHOR CONTRIBUTIONS

The histone modification studies were designed by CJP and IM, performed by CJP and LAF, and analyzed by LTG and CJP. The SETD7 molecular and behavioral studies were designed by JAB and CJP, performed by JAB, LTG, SNB, and CJP, and analyzed by JAB, LTG, and CJP. Bioinformatic analysis was performed by ASC, MT, JAB, and CJP. The physiology experiments were designed by MCC with input from CJP, performed by MCC, MRB, JMT, and LZF, and analyzed by MCC and MRB. The manuscript was written by CJP, LTG, JAB, and MCC, with input from all authors.

## METHODS

### Mice

C57BL/6J mice were maintained on a 12 h light/dark cycleC57BL/6J mice were maintained on a 12 h light/dark cycle (lights on at 4 am) with *ad libitum* access to food and water. All experiments were conducted in accordance with the guidelines of the Institutional Animal Care and Use Committee at Mount Sinai (histone modification study), Princeton University (molecular and *Setd7* RNA-seq and behavior studies), and Washington University in St. Louis (electrophysiology study). Virgin males and females were obtained (Jackson) and bred in-house in trios, males removed after 3-5 days, and females separated to individual cages 20 days after breeding.

#### Early life stress (ELS)

Litters were randomly assigned at birth to standard-facility rearing (Std) or early life stress (ELS) conditions. ELS was performed from postnatal day P10-17 as previously described (*5–7*). Briefly, ELS consisted of 3-4 hours of maternal separation at random times during the light cycle wherein all pups from a litter were removed to a clean holding cage and in addition, EnviroDri nesting material in the home cage was reduced to approximately ½ of standard-reared cages during the days of separations and then restored. Pups were weaned at P21 into cages by litter and sex and left undisturbed except for regular cage changes and checks. Adult male mice were used for the histone profiling experiment. Both males and females were included in *Setd7* experiments. Retired Swiss-Webster male breeder mice (Taconic) were housed individually and used as aggressor mice in chronic non-discriminatory social defeat stress.

#### Transgene expression in juvenile mice

Standard-reared male and female pups were randomly assigned to either experimental *Setd7*-overexpression group or a control GFP group. At P14 (behavior), P21-24 (electrophysiology), or P28 (protein validation), C57BL6/J mice were anesthetized with isoflurane (2-3% induction; 1-3% maintenance) for surgeries and fitted into dual-arm stereotax. Pups were installed with the nose fitted in the nose cone above the bite bar, and the head stabilized using blunt ends of ear bars between the ear and jaw. To bilaterally target the VTA, two Hamilton syringes were angled inward 7°, the Bregma-Lambda distance was measured, the Lambda was taken as the starting point. The following coordinates were used: ML: ± 0.7 mm, AP: +0.86 mm from Lambda, DV: -4.2 mm (P14), or ML: ± 1 mm, AP: +0.91 mm from Lambda, DV: -4.6 mm (P21-28). 300 nL of AAV (1.9 × 10^13^) were injected on each side for behavioral and electrophysiology experiments and 500 nL for protein validation. Both VetBond glue and two sterile sutures were used to ensure complete healing of incisions. Mice were given a perioperative topical dose of bupivacaine (0.25%, 2 mg/kg) and an additional dose 24h following surgery. Mice recovered in a partially-warmed cage before being returned to their home cage. To prevent cannibalization of P14 pups by dams following surgery, dams were habituated with a piece of paper towel with sterile eye lubricant (Puralube Vet Ointment Sterile Ocular Lubricant, Dechra) and liquid tissue adhesive (VetBond, 3M); pups were only returned to the home cage with the dam once completely ambulatory. No qualitative differences in coat quality, grooming, gross locomotor behavior, or water drinking were detectable during testing following surgery. Mice were then aged to adulthood (P60-70) in standard housing conditions.

#### Adult Stress

##### Chronic Non-Discriminatory Social Defeat Stress (behavior, RNA-seq experiments)

In order to test whether epigenetic priming by SETD7-driven H3K4me1 enrichment enhances stress sensitivity, male and female mice were exposed to either control conditions or mild “non-discriminatory” social defeat stress (“defeat”) during the light phase (*49*), as previously described (*56*). Briefly, male and female mice experienced daily social defeat by a Swiss Webster retired breeder (“aggressor;” Taconic) together for 5 days. Female mice were then tested for behavior while male mice experienced 5 more days of social defeat, without a female present, based on previous work documenting that as few as 4 days of adult stress in females is sufficient to induce similar behavioral changes as chronic stress in males (*70, 71*). On each day of social defeat, an experimental male mouse was introduced to an aggressor’s cage (standard rat cage filled with corncob bedding) first for ∼3 minutes, then an experimental female mouse was added into the cage for an additional 5 minutes. Males were then moved across a perforated plexiglass barrier within the aggressor cage, while females were single-housed in a separate rat cage, with a handful of soiled aggressor bedding. Experimental mice were introduced to a new aggressor each day, and males and females rotated in different directions so that trios were unique each day. However, aggressor mice in these experiments were observed to attack less overall on days when females were present, and thus the adult stressor was considered mild for these experiments compared to the standard ten days of male-only social defeat stress. Control mice were housed in a standard mouse cage in pairs of the same sex, separated by a perforated plexiglass barrier.

##### Acute Variable Stress (electrophysiology experiments)

Mice for electrophysiology experiments were subject to three days of variable stress, identical to that previously performed in (*6*) shown to effectively produce behavioral alterations among early-life stress-exposed female mice. Briefly, across three consecutive days, male and female mice were subject to one hour of stressor per day, consisting of either tail suspension, restraint, or mild foot shocks (100 random mild foot shocks at 0.45 mA for 1 h). Electrophysiological recordings were performed on the fourth day.

### Behavioral testing

#### Social avoidance

One day after the last day of social defeat stress, mice were assessed for social avoidance in a social interaction test as previously described (*5*). Social avoidance has been associated with other depression-like behaviors and is responsive to antidepressant treatment (*10, 51*). The test was performed in two stages: during the first 2.5 min, each experimental mouse freely explored an arena (44 × 44 × 20 cm) containing an enclosure centered against one wall of the arena made of plexiglass and wire mesh (10 × 6 cm). The experimental mouse was then returned to its homecage while a novel Swiss-Webster mouse (aggressive retired male breeder) was placed into the enclosure. The experimental mouse was then immediately returned to the arena and allowed to explore the arena freely for 2.5 min. Time spent in the ‘interaction zone’ (14 × 26 cm) surrounding the enclosure, ‘corner zones’ (10 × 10 cm), and ‘distance traveled’ within the arena was measured by video tracking software (Ethovision, Noldus). A social interaction ratio (SI Ratio) was calculated as time spent exploring the novel mouse over time exploring the novel enclosure object. Mice were considered “susceptible” to social defeat if SI Ratio <0.8, “resilient” if SI Ratio >1.2, and “indifferent” for interaction scores in between.

#### Open-field test

Exploratory and anxiety-like behavior was measured by open field exploration one day after social interaction testing. For 10 minutes, mice were allowed to freely explore an empty arena (44 × 44 × 20 cm). Total distance traveled, velocity and time spent in the center (20 × 20 cm) were recorded and measured by video tracking software (Ethovision, Noldus), as previously described (*5*).

### Generation of a Setd7-overexpression viral vector

To directly test the hypothesis that elevated levels of H3K4me1 sensitized mice to future stress experience, we overexpressed the monomethyltransferase *Setd7* in developing VTA. To test this hypothesis, we first generated an adeno-associated virus (AAV8) carrying a *Gfp*- and Myc-tagged *Setd7* transgene (pAAV-EF1a-EGFP-P2A-Myc-Setd7-WPRE-hGHpA). cDNA for *Setd7* (#87131; originally gifted from Francesca Spagnoli; (*72*)) and *Gfp* (pAAV-EF1a-EGFP-WPRE-hGHpA) were purchased from Addgene (#60058; originally gifted from Brandon Harvey; (*73*)). AAV-*Setd7* and AAV-*Gfp* control (identical, without *Myc-Setd7*) vectors were cloned and packaged by the Princeton Neuroscience Viral Core.

### Histological and Molecular Methods

#### Tissue collection for histone modification profiling and RNA extraction

Brains were harvested from adult mice by cervical dislocation and rapid removal into ice-cold PBS. Using an ice-cold slice matrix, bilateral 1 mm-thick 16-gauge punches of VTA were taken fresh and flash-frozen in tubes on dry ice. For mass spectrometry, bilateral VTA tissue punches from three male mice were pooled into a single tube for each sample for processing. Pups from the same litter were combined whenever possible, and pups from different litters of the same condition (Std vs ELS) were pooled otherwise. For RNA extraction (qPCR or sequencing), each sample consisted of tissue from a single male or female mouse.

#### Histone purification and bottom-up mass spectrometry

Three pooled samples were used from each sample per condition (n=3 each for Std and ELS, no adult stress). Extraction of histones and preparation of MS-ready samples were performed as described (*74*). Briefly, histones were extracted in acid from isolated chromatin and purified by TCA precipitation and acetone wash. The purified histones were estimated using Bradford assay and approximately 10-20ug was used to derivatize unmodified lysines, in a mixture of propionic anhydride and acetonitrile (1:3 ratio v/v) for 20 minutes at room temperature. This was followed by digestion with 1µg trypsin/20 ug of histones, 6 hours to overnight at room temperature. Subsequently, derivatization was repeated to propionylate all free N-termini of the peptides. These samples were desalted prior to LC-MS analysis using C18 Stage-tips. Peptide concentration was measured using BCA assay to inject same amount of sample peptides. Quantification of histone peptides were performed in nanoLC coupled online to an LTQ-Orbitrap Elite mass spectrometer (Thermo Scientific). About 1-5 µg of desalted samples were then separated using a 75 µm ID x 17 cm Reprosil-Pur C18-AQ (3 µm; Dr. Maisch GmbH, Germany) nano-column mounted on an EASY-nLC nanoHPLC (Thermo Scientific, San Jose, Ca, USA) in a gradient of 2% to 28% solvent B (A = 0.1% formic acid; B = 95% MeCN, 0.1% formic acid) over 45 minutes, from 28% to 80% solvent B in 5 minutes, 80% B for 10 minutes at a flow-rate of 300 nL/min. and data were acquired using data-independent acquisition (DIA). Briefly, full scan MS (*m/z* 300−1100) was acquired in the Orbitrap with a resolution of 120,000 (at 200 m/z) and an AGC target of 5×10e5. MS/MS was performed in the ion trap with sequential isolation windows of 50 *m/z* with an AGC target of 3×10e4, a CID collision energy of 35 and a maximum injection time of 50 msec. MS/MS data were collected in centroid mode.

### Statistical analysis for mass-spec

For quantification, total area under the extracted ion chromatograms (AUC) were considered for each PTM-bearing peptide and its unmodified form. This was expressed as relative ratio (AUC of any form of the peptide to sum of AUCs of all forms of the peptides with the same amino acid sequence, both modified and unmodified). For isobaric peptides, the relative ratio of two isobaric forms was estimated by averaging the ratio for each fragment ion with different mass between the two species. Within each sample and for each peptide fragment, the proportion of unmodified or modified peptide was calculated by summing mass-spec areas of all modified peptides in that fragment and calculating simple ratios from the total.

Alterations in histone PTHM proportions at each peptide residue were visualized as parts of a whole given their proportional dependency. Given the compositional nature of the data, we first applied an Arcsin transformation to mitigate heterogeneity and constant sum constraints which introduces interdependence between variables (*75*). Arcsin transformations convert proportions between 0 and 1 into values also bounded by 0 and 1, and performed better with “0” data (present in the mass-spec) than log transformations which are also commonly used with compositional data. We then performed two-way repeated measures ANOVAs to specifically assess interactions between rearing condition and proportion of modifications. Two-tailed independent Student’s t-tests were used to assess differences in proportion at each histone modification between Std and ELS groups, as previously reported (*24, 74*). Cohen’s *d* was also calculated to determine effect size, with |*d*|>0.8 considered a large effect size. For visualization and interpretation purposes, log_2_(Fold-Change) in proportion was calculated from control samples for each modification, with which post-hoc tests and effect sizes are reported. All significance thresholds were set at *p*<0.05, with *p*<0.1 also reported for large effect sizes.

### Western Blotting

To validate our viral construct’s ability to increase SETD7 protein and the post-translational modification H3K4me1, and assess whether other related histone modifications were also altered by *Setd7* over-expression, we performed relative protein quantifications via Western blotting. A cohort of wild-type mice were injected with AAV-*Setd7* or AAV-*Gfp* in the VTA at P28. Fourteen days after surgery, 32 male mice were rapidly cervically dislocated, brains were removed immediately, placed into ice-cold PBS, and sliced into 1 mm-thick coronal sections in a brain matrix. Bilateral punches were made from VTA (16-gauge) and were homogenized on ice, in hypotonic lysis buffer (10 mM Tris, 1 mM KCl, 1.5 mM MgCl2, pH 8.0) supplemented with 1 mM Dithiothreitol (DTT), protease inhibitor cocktail (cOmplete™, EDTA-free, Millipore-Sigma) and histone deacetylase (HDAC) inhibitor (10 mM sodium butyrate). Samples were centrifuged for 10 minutes at 10,000xg at 4°C to extract nuclei. Supernatants were discarded and pellets were resuspended in RIPA buffer (50 mM Tris, 150 mM sodium chloride, Triton X-100, 0,1% SDS, pH 8.0) supplemented with 10 mM DTT, protease inhibitor cocktail (cOmplete™, EDTA-free, Millipore-Sigma) and HDAC inhibitor (10 mM sodium butyrate). Protein concentration was evaluated with a 660nm Protein Assay (Pierce). Proteins were separated by SDS-PAGE gels (4-20%, BioRad) and transferred to 0.2 µm low-fluorescence PVDF membranes (Amersham Hybond). The following antibodies were used: H3 (1:5,000, ab24834, Abcam), H3K4me1 (1:2,000, ab8895, Abcam), H3K4me3 (1:1,000, ab8580, Abcam), H3K27ac (1:1,000, 07-360, Millipore-Sigma), SETD7(1:500, TA503322, Origene), and GAPDH (1:2,000, 2118, Cell Signaling). Secondary antibodies were Alexa Fluor 647 anti-rabbit (1:1,000, 711-605-152, Jackson ImmunoResearch) and AffinipureCy3 anti-mouse (1:1000, 715-165-150, Jackson ImmunoResearch). Images were acquired with Azure Imaging System (Azure Biosystems) and quantified using ImageJ.

### RNA extraction

On the day following behavioral testing, mice were sacrificed by rapid cervical dislocation. Brains were rapidly removed, and bilateral 1 mm-thick 16-gauge punches of VTA were taken fresh, flash-frozen on dry ice, and stored at -80C until processing. RNA was extracted from tissue samples with Trizol (Invitrogen) and chloroform (Sigma) and purified with RNeasy Micro Kits (Qiagen). Separate cohorts of mice were used for qPCR and sequencing experiments. For qPCR experiments, tissue and cDNA was used from a previous study (*76*).

### qPCR

For relative quantification of histone methyltransferases in VTA after ELS, cDNA was created with a reverse-transcription kit (HighCapRT, Applied Biosystems). Primers for real-time semi-quantitative PCR (qPCR) were validated for efficiency and specificity. The following primer pairs were used: *Setd7* (Forward: GCAGGGCCAGGGCGTTTACA; Reverse: GGCCGGTCCATTCAGCTCTCC); *Kmt2a* (also known as MLL1; Forward: TGCTAAAGGGAACTTCTGCCC; Reverse: TCTCATCTTCTGTACCTGAGAGACT); *Hprt* (as housekeeping gene; Forward: GCAGTACAGCCCCAAAATGG; Reverse: GGTCCTTTTCACCAGCAAGCT). All qPCR reactions were run in triplicate and used SYBR-green on a QuantStudio 7 (ThermoFisher). The 2^(-ΔΔCt) method was used to calculate RNA expression analysis compared to the standard-reared group (including both males and females) with *Hprt* as a control reference gene (*77*). In addition, we report normalized reads (reads per thousand-mapped, RPKM) of published RNA-seq data from male VTA [Gene Expression Omnibus (GEO) accession GSE89692; (*5*)] for histone-modifying genes of interest.

### RNA-sequencing

We analyzed changes in gene expression across the transcriptome by RNA-sequencing following viral-mediated *Setd7* or *Gfp* expression, social defeat, and behavioral testing. RNA was converted into libraries for RNA-sequencing using an Illumina TruSeq RiboZero kit. Samples each reflect one individual mouse, except for one sample in which two female *Setd7*-CNSDS mice were combined due to insufficient RNA concentration. After confirming the quality of the cDNA library, it was sequenced for paired-end reads to a read depth of over 15 million (15M) reads per sample at the Lewis-Sigler Institute for Integrative Genomics (Princeton University)

### RNA-seq data analysis

Reads from the Illumina TruSeq RNA library were analyzed using the Galaxy Genomics platform. Quality checks were performed using FastQC. Reads were then mapped to a reference mouse genome (GRCm39) using Bowtie2. Using the FeatureCounts tool, the mapped reads were counted across mouse genes and assessed for gene body coverage to check for potential bias in the quantity or quality of reads along the genome. DESeq2 was used to normalize read counts and perform differential gene expression (DEG) analysis with standard parameters and filtering. Principal component analysis by DESeq2 identified one male *Setd7*-CNSDS mouse sample as an outlier, and the sample was removed from further analysis. Two additional samples were removed for having 1.5 SD lower *Scl6a3* and *Th* CPM, indicating tissue punching error that included less VTA than other samples. Thus, our final DEG analysis included 3 GFP-Ctrl, 4 Setd7-Ctrl, 4 GFP-SD, and 2 Setd7-SD samples. Males and females were included in each group and combined for group-level analysis. DEGs were defined as genes with *P*.*adj*<0.1 for gene ontology enrichment analysis and calculating percentage of up- and down-regulated genes. In volcano plots, genes with *P*.*adj*<0.05 are shown in darker colors and additional genes with uncorrected *p*<0.01 and Log_2_FC<|0.375| shown in lighter colors. Gene ontology enrichment analysis was performed using the database for annotation, visualization, and integrated discovery (DAVID) online tool with a minimum count of 4, displaying only non-redundant terms. Two-sided rank-rank hypergeometric overlap analysis was done as previously described (*6, 78, 79*) using genes with baseMean expression>10.

### Immunohistochemistry and imaging

A portion of the mice that received viral overexpression and underwent behavioral testing were used to assess targeting and specificity of viral expression. Mice were sacrificed a day after the last behavior testing. Mice were anesthetized with ketamine/xylazine (100 mg/kg and 10 mg/kg, i.p.) and transcardially perfused with sterile 1X PBS followed by 4% paraformaldehyde. Brains were removed, equilibrated in 30% sucrose in PBS, and frozen at -80°C until processing. Brains were sectioned at 50 µm-thick slices using a cryostat. Immunohistochemistry was performed using the following antibodies: SETD7 (Origene, TA503322), GFP (Aves Lab, GFP-1020), and Prolong Gold Antifade with DAPI (Invitrogen). Imaging was performed on Hamamatsu NDP slide scanner (Hamamatsu Nanozoomer 2.0HT).

### Patch-clamp electrophysiology

Mice were injected with AAV-*Gfp* or AAV-*Setd7* from P21-24 and reared normally until adulthood. Mice in the control group (n=5/4 male/female each for *Gfp* and *Setd7*) were sacrificed directly from the home cage and brains were removed and sliced horizontally through the VTA on a vibratome at 200 µm in ASCF as in (*80, 81*)). Mice in the stress group (n=5/4 male/female each for *Gfp* and *Setd7*) underwent three days of variable stress (described above; (*6*)) and recordings were made the day after the last stressor.

Patch-clamp electrophysiology data acquisition was completed using Clampex 11.4 software (Molecular Devices). Visualized whole-cell recording techniques were used to measure holding and synaptic responses of DA neurons in the VTA. Currents were amplified, low-pass filtered at 4 kHz, and digitized at 10 kHz. Recordings were performed in whole cell configuration with the patch pipette internal solution containing (in mM): 130 potassium gluconate, 10 phosphocreatine disodium salt, 4 MgCl_2_, 3.4 Na_2_ATP, 0.1 Na_3_GTP, 1.1 EGTA, 5 HEPES, with a pipette resistance between 2.8 and 3.2 MΩ. All recordings were performed in dopamine neurons identified by a cellular capacitance > 30 pF, spontaneous firing below 5 Hz, and the presence of a hyperpolarization-activated inward current, /_h_ measured in voltage-clamp configuration. To further ascertain the presence of /_h_ currents, neurons were voltage clamped at −70 mV, and the response to five hyperpolarizing voltage steps (250 ms, −25 to −125 mV in increments of −25 pA) were measured. Input resistance and the responses to current injections were made in current-clamp configuration. Input resistance was calculated following the average membrane change in response to 20 successive injections of -10 pA of current. To generate an input-output curve of neuronal excitability, 2000 ms current steps were injected in increments of 20 pA from −60 to 200 pA and the numbers of spikes were quantified for each step. Voltage sag was measured by injecting negative current steps in -10 pA increments until a resting membrane voltage of -100 pA was reached. Action potential threshold was determined by repeatedly injecting 2000 ms ramps of increasing current (0 — 200 pA) for 5 trials and calculating the mean voltage threshold of the first action potential. Recordings were excluded if the series resistance or noise level varied by more than 20% over the course of recordings.

### Statistical analysis for gene expression, protein quantification, and behavior

Statistical analyses were performed using SPSS (Version 26), Prism (GraphPad; version 9) or Statsmodels (Python 3.12), with alpha set to 0.05. Outliers were defined as more than two standard-deviations away from group mean within cohort and were removed to avoid type-two error. Effects of ELS alone were assessed by Student’s t test and Cohen’s *d* effect sizes were calculated as *d*=(meanB-meanA)/(total StDev). |*d*|>0.8 is considered a large effect size. Effects of early-life and adult stress together were assessed using ANOVA or mixed-effects modeling, as indicated. Post-hoc analyses between pairs of groups were conducted using Sidak’s multiple-comparisons correction if there were significant main effects or interactions. Significant changes in proportion of susceptible or resilient mice in the social avoidance task were calculated by two-tailed two-proportion z-test. All data are plotted as mean ± the standard error of the mean (SEM).

## Data Availability

All relevant data that support the findings of this study are available by request from the corresponding author (CJP). RNA-sequencing data will be deposited with GEO.

---

**Supplemental Figure 1:**
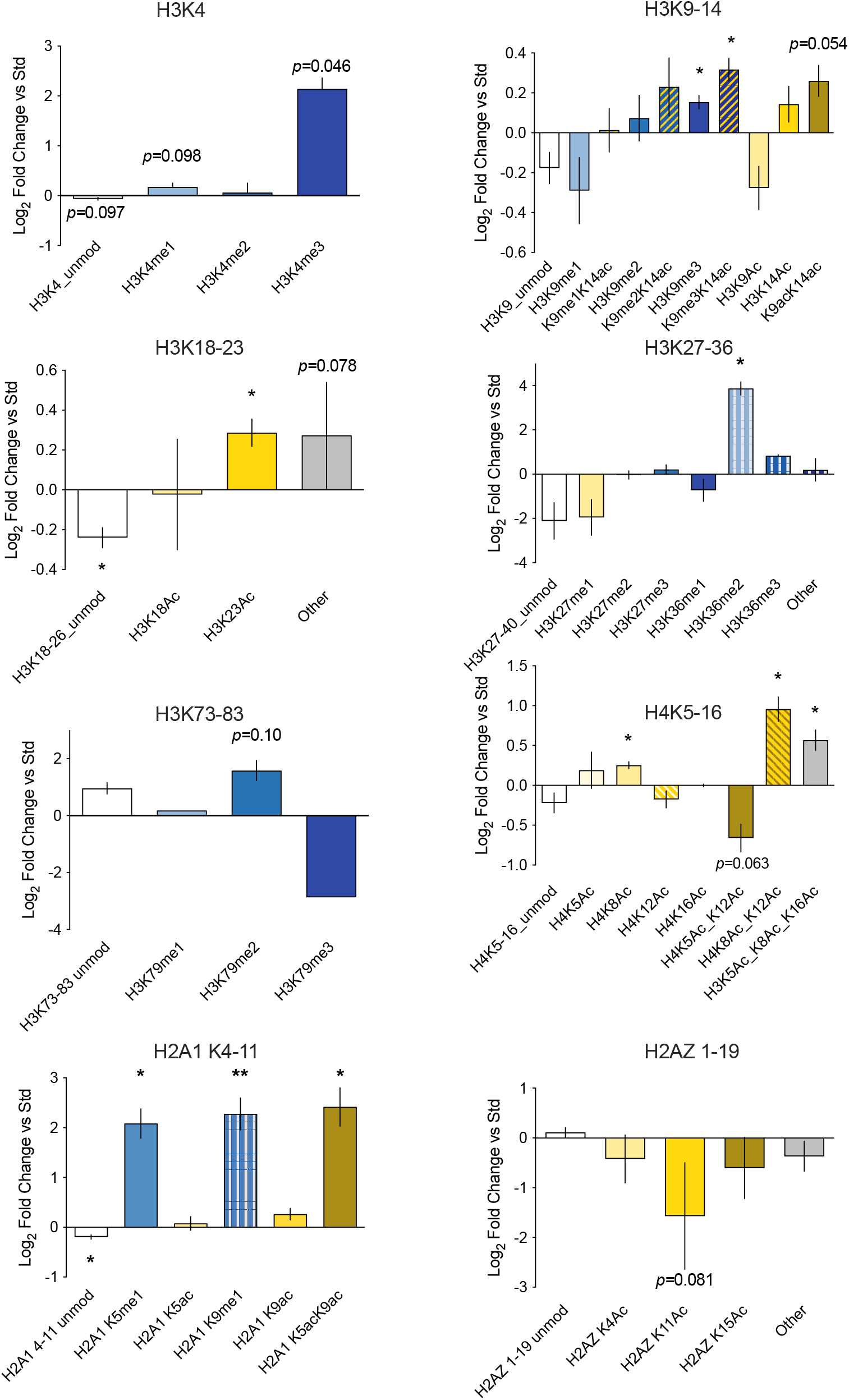
Pairwise Change in PTHMs in the VTA following ELS. Log_2_ fold-change of PTHM’s within each histone fragment, normalized to Std, and represented as mean ± SEM. Within PTHM fragments, ELS increased proportions of the following PTHMs/combinations with large effect size (Cohen’s |*d*|>1.3): H3K4me1 (*d*=1.337; *p=*0.098), H3K4me3 (*d*=1.495; *p=*0.046), H3K9me3 (*d*=1.560; *p=*0.031), H3K9me3-K14Ac (*d*=1.554; *p=*0.032), H3K9Ac-K14Ac (*d*=1.468; *p=*0.054), H3K23Ac (*d*=1.506; *p=*0.043), H3K36me2 (*d*=1.608; *p=*0.020), H4K8Ac (*d*=1.616; *p=*0.019), H4K8Ac-K12Ac (*d*=;1.687 *p=*0.008), H4K5Ac-K8Ac-K16Ac (*d*=1.625; *p=*0.018), H2A.1K5me1 (*d*=1.607; *p=*0.21), H2A.1K9me1 (*d*=1.687; *p=*0. 009), and H2A.1K5Ac-K9Ac (*d*=1.505; *p=*0.043). ELS decreased proportions of: unmodified H3K3-8 (*d*=1.341; *p=*0.095), unmodified H3K18-26 (*d*=1.630; *p=*0.017), unmodified H2A.1K4-11 (*d*=;1.669 *p=*0.011), H4K5Ac-K12Ac (*d*=1.436; *p=*0.063), and H2AZK11Ac (*d*=1.384; *p=*0.081). Statistics are two-tailed t-tests on arcsin-transformed data, represented by \**p*<0.05; *p* values <0.1 are shown.

